# Programming CRISPRi-dCas9 to control the lifecycle of bacteriophage T7

**DOI:** 10.1101/2024.05.15.594216

**Authors:** Tobias Bergmiller

**Affiliations:** University of Exeter, Biosciences, Stocker Road, EX4 4QD Exeter, United Kingdom

## Abstract

CRISPR interference (CRISPRi) based on catalytically dead Cas9 nuclease of *Streptococcus pyogenes* is a programmable and highly flexible tool to interrogate gene function and essentiality in bacteria due to its ability to block transcription elongation at nearly any desired DNA target. Here, I assess how CRISPRi-dCas9 can be programmed to control the life cycle and infectivity of *Escherichia coli* bacteriophage T7, a highly virulent and obligatory lytic phage, by blocking critical host-dependent promoters that are required for host cell entry and life cycle execution. Using fluorescent reporters demonstrates that the efficacy of CRISPRi-dCas9 towards *E. coli* RNAP promoters depends on target promoter strength, and that the phage’s own T7 RNAP can be downregulated very efficiently. Effects on the time to lysis were partially dictated by the chronological order of phage DNA and dCas9 target appearance in the cell, suggesting an accessory role of *E. coli* RNAP during the early stages of T7 genome ejection. The stringency of the CRISPRi-dCas9 approach was greatly improved when using multiplex sgRNAs to target multiple phage regions simultaneously, resulting in a 25% increase in the time to lysis and up to 8-fold reduction in plaque size. Overall, this work expands CRISPRi-dCas9 as a flexible tool to non-invasively manipulate and probe the lifecycle and infectivity of otherwise native T7 phage.

## Introduction

Clustered Regularly Interspaced Short Palindromic Repeats interference (CRISPRi) is a powerful tool to study genotype-phenotype associations or bacteriophage (phage)-host interactions in bacteria (Qi et al., 2013; Peters et al., 2016; Peters et al., 2019; Mutalik et al., 2020; Rishi et al., 2020), and to engineer *de-novo* synthetic gene regulatory circuits and control (Kuo et al., 2020; Rueff et al., 2023). A commonly used CRISPRi approach stems from the CRISPR-Cas9 system of *Streptococcus pyogenes* and utilises a catalytically dead Cas9 (dCas9) nuclease that doesn’t cleave DNA but sterically blocks transcription elongation by RNA polymerase, leading to repression or knockdown of the target gene (Qi et al., 2013). Transcription blockage requires a small guide RNA (sgRNA) that programs dCas9 to bind to the non-coding strand at a specific DNA target containing a NGG protospacer-adjacent motive (PAM). Virtually any gene or non-coding region containing a PAM can thus be targeted by dCas9 and its expression level manipulated with very high specificity (Qi et al., 2013; Rousset et al., 2018; Rishi et al., 2020).

In bacteria, this approach is titratable and requires only minimal engineering of the target strain by introducing an inducible dCas9 gene and sgRNA (Rousset et al., 2018; Rishi et al., 2020). The latter is commonly supplied on a low to medium-copy plasmid. This enables straight-forward and cost-efficient investigation of gene function across many bacterial species through the design and construction of sgRNAs or sgRNA libraries for high-throughput studies, replacing the time-consuming construction of gene knockouts or transposon mutant libraires (Peters et al., 2019). Also, expression of more than one sgRNA enables multiplex approaches to manipulate several genetic targets at once, to explore combinatorial effects (Ellis et al., 2023), and to engineer metabolic pathways and flux (Kim et al., 2017). Other engineering avenues have explored using CRISPRi-dCas9 in bacteria to activate transcription by fusing transcriptional activation domains to dCas9 (Dong et al., 2018), or to construct more complex regulatory and repressilator-like circuits (Kuo et al., 2020; Rueff et al., 2023). Until recently, CRISPRi-dCas9 has only been used to study and manipulate the genomic content or plasmids in bacteria, but not to target bacteriophage (phage) genomes to study phage gene function or to control their lifecycles. Phages are viruses that exclusively predate and kill bacteria with high specificity and efficacy, and they are of great interest for therapeutic applications to combat infections caused by multi-drug resistant bacterial pathogens (Strathdee et al., 2023). Of great therapeutic interest are obligatory lytic phages, such as *E. coli* phage T7, that lack the biphasic lysis-lysogeny life cycle of temperate phages, and that eliminate their bacterial hosts with high specificity and efficacy. T7 is a highly virulent phage that has been extensively studied over the last decades (Calendar, 2006). Its host range can be synthetically expanded to other Gram-negative species (Yehl et al., 2019), and its small 40kb genome can be reassembled using standard molecular techniques and rebooted *in-vivo* or *in-vitro* using cell-free systems (Levrier et al., 2024). This versatility makes T7 easy to manipulate and of great interest for therapeutic and biotechnological applications.

T7 phage has a highly modular 40kb genome separated into three modules: the early region including an internalisation signal and genes required to hijack the host and T7 RNAP, DNA replication, and phage assembly. These modules are arranged in the chronological order of transcription events that take place during genome entry (Calendar, 2006). T7 phage ejects about 850bp of its genome into the host cell, and further genome internalisation stringently requires catalysis by the hosts RNA polymerase (RNAP) (Zavriev and Shemyakin, 1982; Garcia and Molineux, 1995). This is facilitated by an array of three strong promoters, *A1*, *A2*, and *A3*, that are located on the ejected and otherwise non-coding 850bp “early” region that serves as internalisation signal (Dunn and Studier, 1983). Once further genome entry by *E. coli* RNAP has occurred, expression of the phages own T7 RNAP, initially driven by the hosts RNAP from promoter *C* on the T7 genome, leads to full genome internalisation and lifecycle execution (Garcia and Molineux, 1996). This culminates in host lysis and progeny release 11 minutes after DNA injection at 37°C.

In general, there is limited experimental evidence of the efficacy of *S. pyogenes* CRISPRi-dCas9 towards phages. A report by Bickard et al suggests that the genome of the temperate phage lambda can be targeted by CRISPRi-dCas9 (Rousset et al., 2018), and *in-vitro* studies of T7 RNAP show high efficacy of dCas9 blocking transcription elongation of T7 RNAP (Widom et al., 2019). Recent reports demonstrated the use of dCas12a for functional studies of temperate phages Lambda and P1 in *E. coli* (Piya et al., 2023), and CRISPRi-ART based on dRfxCas13d for a range of different phages (Adler et al., 2023). However, the usability of the popular *S. pyogenes* CRISPRi-dCas9 system to manipulate the lifecycle or infectivity of T7 phage for synthetic biology approaches such as to engineer an infectivity switch (Chitboonthavisuk et al., 2022) or functional studies remain largely unexplored.

In this study, I design, test, and quantify the efficacy of CRISPRi-dCas9 to tune and control the T7 phage life cycle and infectivity by targeting important host RNAP-dependent promoters. I focus on the promoters in the early non-coding region comprising the internalisation signal and the host promoter *C* controlling initial expression of T7 RNAP. I build a set of tools to quantify the efficacy of CRISPRi-dCas9 and to tune the lifecycle of T7 phage. This work is a proof-of concept study that highlights the use of the popular CRISPRi-dCas9 tool to tune the infectivity of lytic phages without the requirement of phage engineering.

## Results

### sgRNA selection and construction

Following T7 phage genome annotations and prior knowledge of the T7 phage lifecycle (Chan et al., 2005), I selected the reportedly strongest and most critical *E. coli* RNAP-dependent promoters *A1*, *A2*, and *A3*, comprising the non-coding internalisation signal (*INTS* hereafter), and the T7 RNAP promoter *C*, as main targets for CRISPRi-dCas9 (**Figure 1 C**). The rationale was that blocking these promoters and gene products interferes with the T7 phage lifecycle at different stages, allowing to control, or at least tune, its progression. More specifically, the hypothesis was that sterically blocking *INTS* may slow down or even abolish T7 infection by hindering *E. coli* RNAP catalysing T7 genome entry. Furthermore, expression of early genes that are required to overcome host defense systems and to shut off host transcription is driven by the *INTS* promoters. Blocking expression of T7 RNAP was another prime target of this approach and followed the idea that blocking transcription of T7 RNAP may hinder progression of the T7 lifecycle to later stages which are entirely T7 RNAP-dependent.

**Figure 1.**
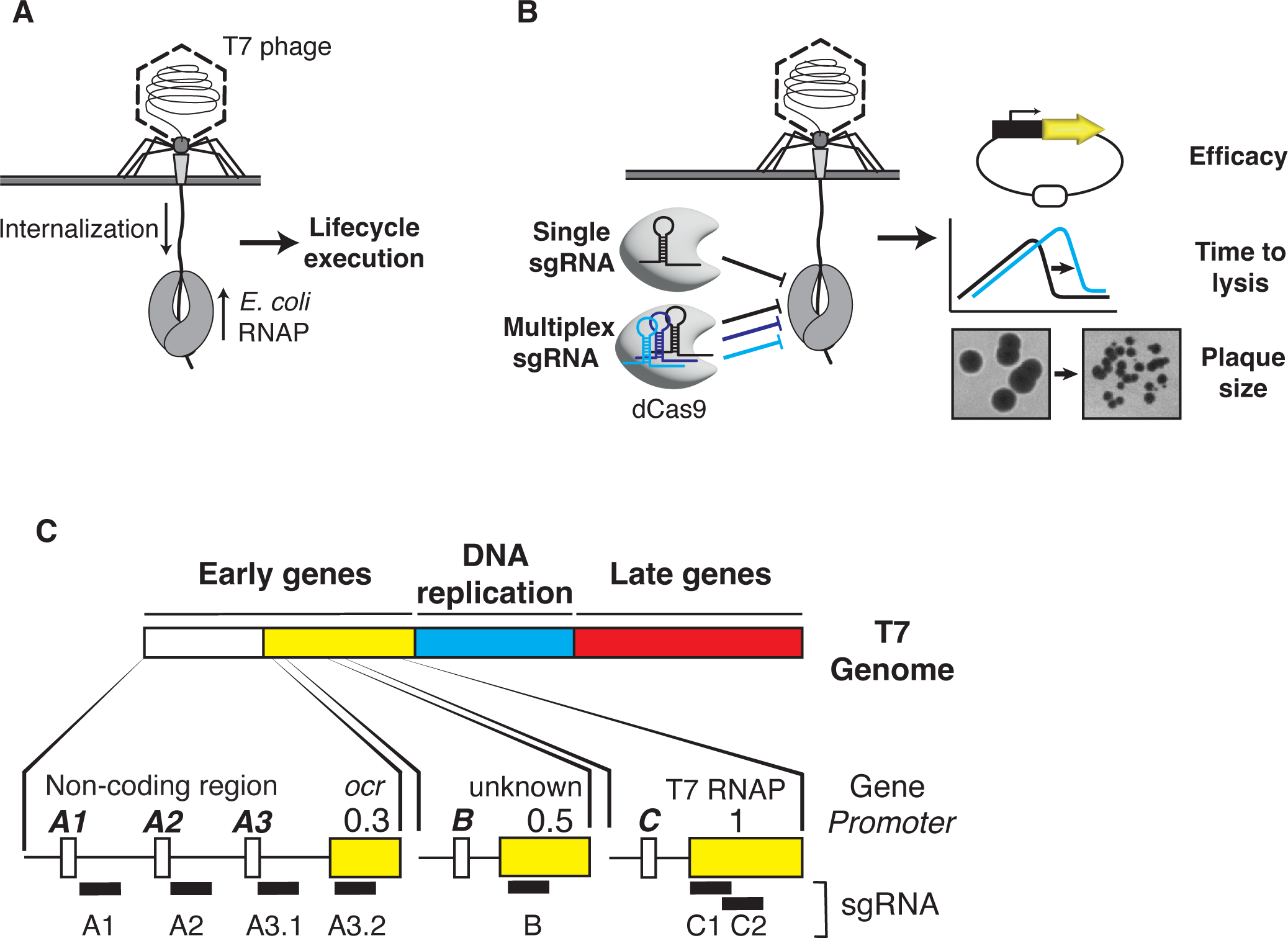
Targeting host RNAP promoters to interfere with the lifecycle of T7 phage. A) Schematic of DNA ejection by T7 phage. T7 ejects about 850bp of its genome into the host cell which comprises the internalisation signal. *E. coli* RNAP binds to the strong promoters on the internalisation signal and catalyses further genome entry and transcription of early genes, leading to coupled internalisation and lifecycle execution. B) Using single or multiplex sgRNAs, dCas9 is programmed to interfere with *E. coli* RNAP at specific sites along the T7 phage genome. Fluorescent reporters are used to assay the efficacy of this approach using fluorescent knockdown experiment, and batch culture and plaquing experiments to quantify the impact of this approach on the time to lysis and infectivity of T7 phage. C) Schematic of the T7 phage genome (not to scale). The lower section shows the targets for the CRISPRi-dCas9 approach in the early region of T7 and approximate positions of sgRNA binding.

I consequently designed a set of sgRNAs using published design principles and where possible, an automated workflow (Banta et al., 2020). The sgRNAs were chosen such that dCas9 homes in on T7 RNAP which is controlled by *C*, and on *INTS* at several positions (**Figure 1C**). sgRNAs towards *INTS* were designed to bind downstream of the annotated transcription start sites of *A1*, *A2*, and *A3* (sgRNA *3.1*), and a fourth sgRNA (sgRNA *3.2*) targeting gp0.3 (*ocr*) within the 5’ end of the open reading frame.

The sgRNA towards *C* was designed to bind within the 5’ end of the T7 RNAP ORF to have maximal knockdown effect on T7 RNAP. sgRNAs were then cloned into a medium copy sgRNA expression plasmid (**see Material and Methods**). I also constructed two multiplex sgRNA arrays consisting of 3 (3x) and 5 (5x) sgRNAs to target *INTS* and *C* at several sites simultaneously. To maximise the effect of the 5x multiplex sgRNA approach, promoter *B* was included as a multiplex target. Promoter *B* can have accessory function for T7 genome internalisation (Garcia and Molineux, 1996) and controls expression of gp0.5 whose function is unknown. In addition, I constructed strain TB549 that harbours a chromosomally inserted single-copy and arabinose-inducible P*_araBAD_*-dCas9 construct (Haldimann and Wanner, 2001). Expression of dCas9 in TB549 by adding arabinose had no measurable effects on growth rate, and constitutive expression of sgRNAs in TB549 in the absence or presence of dCas9 expression also showed no negative effects of growth (**SI Figure 1**).

### Quantifying CRISPRi-dCas9 efficacy using fluorescent reporters

Next, I constructed fluorescent transcriptional reporter plasmids to assay the expression strength of *E. coli* RNAP-dependent promoters encoded by T7 phage, and to quantify the efficacy of transcriptional repression by CRISPRi-dCas9 by fluorescence knockdown measurements. I build reporter plasmids following a previously published approach (Zaslaver et al., 2006) by placing T7 DNA fragments containing the desired promoters flanked by ∼150bp up-and downstream sequences onto a low copy oriSC101 plasmid (Lutz and Bujard, 1997) in front of a yellow fluorescent protein gene (*yfp* Venus). The *yfp* ORF is preceded by a strong ribosome binding site (RBS), and YFP expression levels report on transcript abundance and transcriptional activity of the promoters of interest. I included all annotated *E. coli* RNAP promoters, *B*, *C* and *E*, and a ∼570bp fragment of *INTS* containing promoters *A1*, *A2*, and *A3*, to compare their relative strengths and to validate the target choice.

The reporter plasmids had mild to strong detrimental effects on the growth rates of their host strains, which agrees with previous findings that T7 phage genome fragments cloned into plasmids can be detrimental or even toxic to growth (Studier and Rosenberg, 1981). Especially *INTS* turned out to be highly toxic and showed a heterogeneous small colony phenotype, and an unusual bi-phasic growth pattern (**SI Figure 2**). A large colony that displayed uniform growth patterns while expressing very high levels of YFP was selected at random. This reporter plasmid had a small deletion upstream of the *ocr* fragment (but outside of promoter *A3* and the sgRNA binding site) that was included in the *INTS* construct (**SI Figure 2**). This stabilised promoter plasmid was used for all experiments shown in this work hereafter.

Assaying YFP expression as a proxy of promoter strength enabled a ranking of *E. coli* RNAP promoters. *E* and *B* were the weakest promoters and slightly weaker or on par with an IPTG-induced P_LlacO1_-YFP control plasmid, while promoter *C* and the *INTS* fragment showed a ∼3-fold and ∼10-fold higher promoter strength than P_LlacO1_-YFP (**Figure 2 B**). Promoters *E* and *B* are described as minor *E. coli* promoters (Studier and Rosenberg, 1981) and their role for T7 infection is unclear. *E* and *B* were consequently excluded from further investigation and analysis that was focused on *INTS* and *C*, apart from targeting *B* with the 5x multiplex sgRNA.

**Figure 2.**
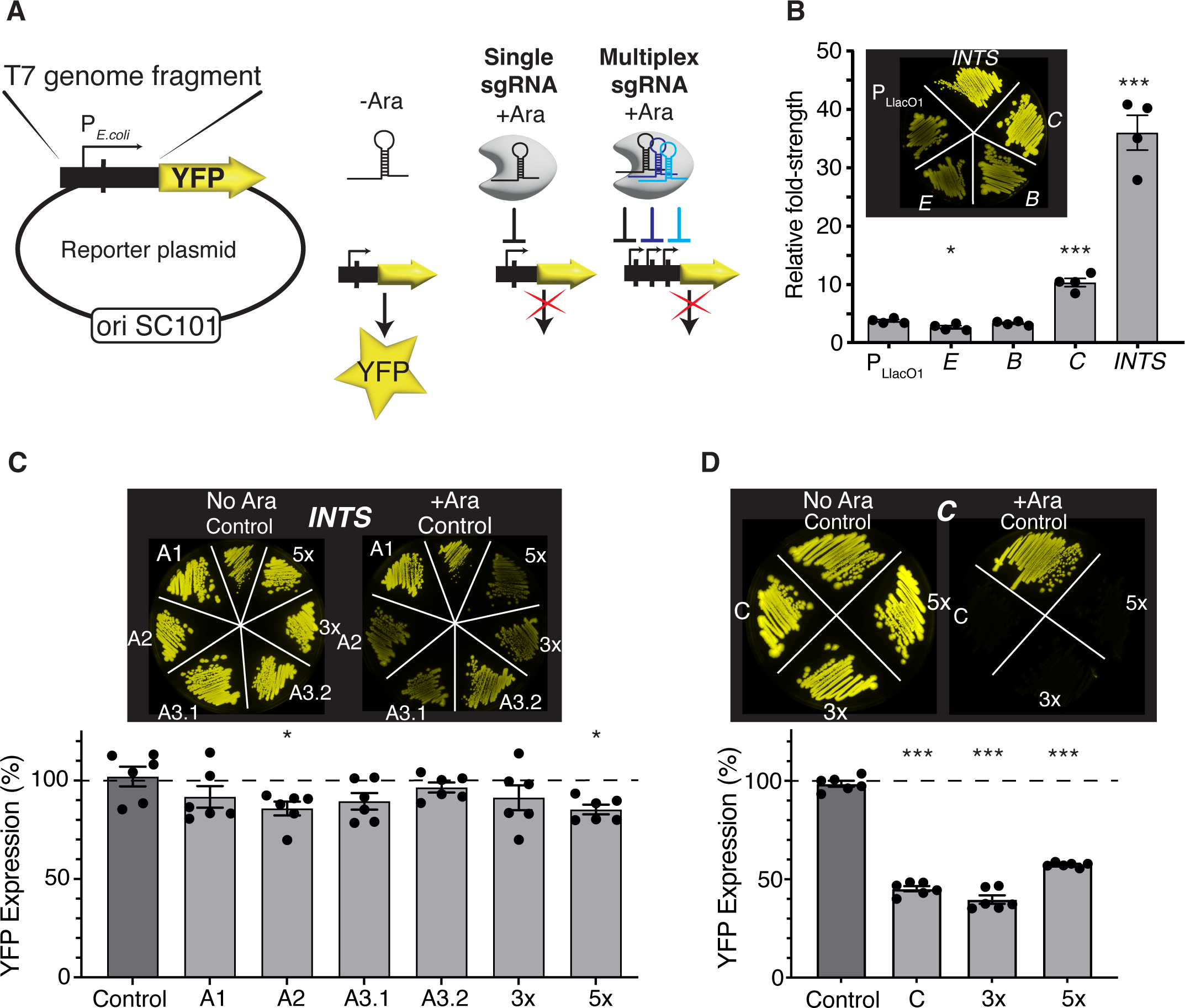
Quantifying the efficacy of CRISPRi-dCas9 using fluorescent reporters. A) Construction of fluorescent *E. coli* RNAP promoter reporter plasmids. T7 genome fragments encoding selected *E. coli* RNAP promoters were cloned into an ori SC101 plasmid (pZS1 (Lutz and Bujard, 1997)) in front of a *yfp* Venus open reading frame which carries its own ribosome binding site (not shown). Transcriptional activation of promoters leads to production of YFP Venus. Targeting promoters with CRISPRi-dCas9 by supplying the inducer L-arabinose and single or multiplex sgRNAs allows to quantify the efficacy of knockdown by measuring YFP fluorescence levels. B) Comparison of *E. coli* RNAP promoter strengths. YFP expression levels were normalised to uninduced pZS12-YFP Venus and shown is fold-change promoter strength. Insert: fluorescence image of TB549 with promoter plasmids and uninduced pZS12-YFP Venus, which is relatively leaky in TB549. p<10^-3^ in a nonparametric t-test comparing induced pZS12-YFP to *C* or *INTS* indicated by ***, and 0.02 for *E* indicated by *. N=4 for all data shown. C) Fluorescence knockdown efficiency across *INTS* or d) *C* (T7 RNAP) using single or 3x and 5x multiplex sgRNAs. The inserts show fluorescence images of TB549 with corresponding sgRNA, and fluorescent reporter plasmids streaked on selective LB agar plates without (left) or with L-arabinose (right). The graphs show fluorescence knockdown efficiency normalised to controls lacking L-arabinose and dCas9 expression. The y-axis shows % increase or decrease in YFP expression. In C), p<0.05 in a pairwise t-test comparing Control sgRNA with A2 or 5x sgRNA indicated by *. For D), p<10^-3^ in a pairwise t-test of C, 3x or 5x to the control sgRNA indicated by ***. N=6 for all data shown. Error bars are one standard error of the mean.

TB549 was then sequentially transformed with sgRNA plasmids and fluorescent reporter plasmids, and first the effects of targeting *A1*, *A2* and *A3* within *INTS* with individual sgRNAs was evaluated. The overall efficacy of CRISPRi-dCas9 using single sgRNAs was low, leading to a ∼15% decrease in YFP expression at best (**Figure 2 C)**. Nevertheless, the order of targeting *A1*, *A2* and *A3* promoters on *INTS* appeared to be important: targeting *A1* or *A3* with an sgRNA binding inside the *ocr* fragment (sgRNA A1 or A3.2) had very weak or no discernible effects on fluorescence, while targeting *A2* and *A3* (sgRNA A2 or A3.1) had mild, and in the case of A2, significant effects (**Figure 2 C**). In contrast, targeting the T7 RNAP reporter driven by *C* was very efficient, leading to a 60% reduction in expression of fluorescence (**Figure 2 D**).

Subsequently, I constructed and tested arrays of 3x and 5x multiplex sgRNAs which contained A3.1, A3.2, and C sgRNAs for the 3x multiplex array, and A2, A3.2, B, and two sgRNAs targeting *C* for the 5x multiplex array. Here, the rationale was that targeting *INTS* at several locations may improve knock-down due to the presence of several roadblocks, or by blocking RNAP access to alternative promoters. Targeting *C* twice and additionally *B* at the same time may facilitate additive even synergistic effects towards blocking the T7 phage lifecycle in later experiments using the same sgRNA constructs. I found that the 5x array showed only a mild but significant improvement in efficacy while the 3x array showed no further improvement in knocking down YFP expression from *INTS* (**Figure 2 C**). Surprisingly, while the 3x sgRNA array was targeting *C* with similar efficacy as the single sgRNA, targeting *C* at two positions simultaneously using the 5x sgRNA had weakly antagonistic effects on YFP knockdown (**Figure 2 D**).

Together, this data suggests that there is no singular dominant expression driver in the *INTS* promoter array: blocking *A1* had no significant effects on YFP expression, likely as *E. coli* RNAP can initiate transcription at promoters *A2* and *A3*. In turn, blocking *A2* or *A3* lowered YFP expression, potentially by limiting the ability of *E. coli* RNAP to initiate transcription downstream at one or multiple alternative promoters. Interestingly, blocking *E. coli* RNAP even further downstream of *A3* within the *ocr* fragment (sgRNA A3.2) had no effect on YFP expression. One possibility is that the very high strength of the promoter array of *INTS* limits the efficacy of CRISPRi-dCas9 to operate as an efficient roadblock of transcription elongation outside of the promoter array. Also, these results show that repeatedly and simultaneously targeting the same region does not necessarily lead to additive or synergistic knockdown effects.

### Efficacy of CRISPRi-dCas9 towards T7 phage during infection

Next, I sought to test the ability of CRISPRi-dCas9 to interfere with the time to lysis of bacterial cultures by T7 phage which can be used as a robust proxy for resilience of the host towards phage infection (Chitboonthavisuk et al., 2022). To do so, I compared the onset of lysis of bacterial cultures expressing sgRNAs and dCas9 with those lacking the dCas9 inducer L-arabinose at identical multiplicities of infection (MOI) and initial bacterial cell densities. I grew replicates of cultures with or without L-arabinose to late exponential phase, diluted them into fresh LB broth and added small amounts of T7 phage (MOI 10^-5^). Addition of L-arabinose and dCas9 expression led to a notable increase of the onset of lysis for cultures expressing A1 and C sgRNA and the 3x and 5x multiplex constructs (**Figure 3A**). The onset of culture lysis was then quantified by determining the inflection point of the growth curve at which population collapse by lysis occurred, and at which the slope turned negative (**Material and Methods**). This showed that the effects of CRISPRi-dCas9 targeting *INTS* followed a slightly different order compared to fluorescent knockdown assays: amongst all individual sgRNAs targeting *INTS*, A1 had the strongest effect on the time to lysis, which increased by ∼22%. The other *INTS* targets followed a similar trend observed in the fluorescent reporter assays: targeting A2 and A3.1 had mild but significant effects while targeting A3.2 at the very end of *INTS* had no effect (**Figure 3B**) on the time to lysis. Overall, this order of effects approximately follows the chronological order of appearance of the *INTS* DNA segment into the cell during T7 DNA ejection, with *A1* appearing first, *A2* second and so on. This means that the efficacy of dCas9 is highest during the very early stages of T7 DNA ejection, which has two interesting consequences for the DNA ejection model: one possibility is that *A1* and upstream sequences are important for RNAP binding and initiation of RNAP-catalysed DNA internalisation, and that dCas9 binding is interfering with this process. The other possibility is that the early T7 DNA eject process is partially dependent on the hosts RNAP, or that RNAP has an accessory role for the early stage of DNA ejection, which is not in line with the experimentally established model of RNAP-independent DNA ejection of T7 phage (Garcia and Molineux, 1995).

**Figure 3.**
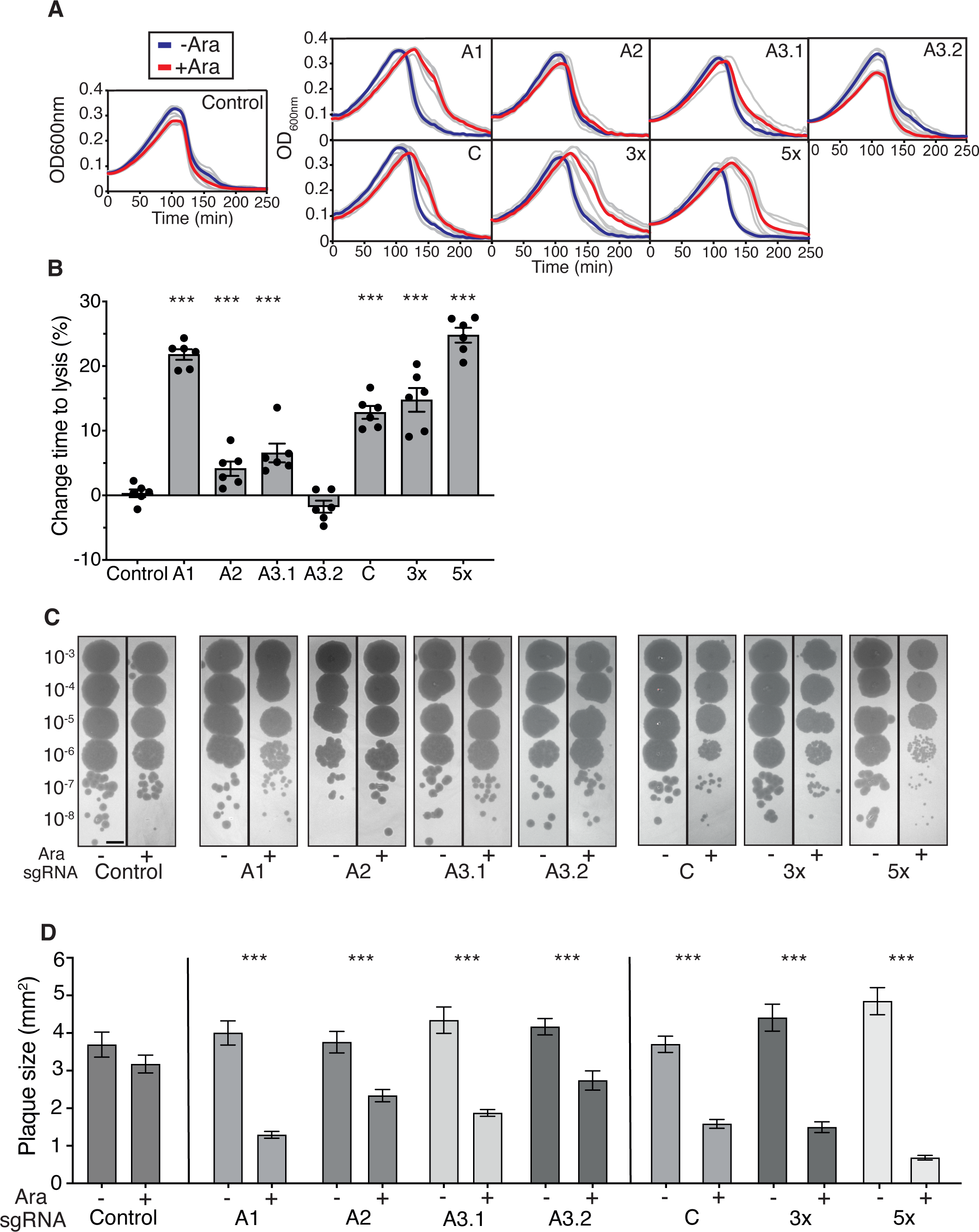
Effects of CRISPRi-dCas9 on the time to lysis, EOP, and plaque size of T7 phage. A) Effects of CRISPRi-dCas9 expression on culture lysis. Shown are growth curves of cultures infected with T7 phage at MOI 10^-5^ with (red) or without (blue) 0.1% L-arabinose. Grey lines are raw data, thick lines are means (n=6). Left, control sgRNA. Top row: individual sgRNAs targeting *INTS*. Bottom row: individual sgRNA targeting *C*, and 3x and 5x multiplex constructs targeting several T7 phage regions simultaneously. B) Quantification of the time to lysis of T7 phage. Onset of culture lysis was quantified by finding the inflection points using a procedure described in Material and Methods. The inflection timepoints of L-arabinose-induced cultures were normalised to their respective controls lacking L-arabinose. Two tailed t-test comparing the Control sgRNA to A3.2 sgRNA shows no significance, all other paired t-tests with Control sgRNA show significance with p<10^-3^ indicated by ***. C) Effects of CRISPRi-dCas9 expression on EOP. Expression of CRISPRi-dCas9 has no observable effect on EOP, but noticeable effects on plaque size. Shown are regions of interest of equal size, scale bar = 5mm. D) Quantification of plaque size shows effects of CRISPRi-dCas9. Shown is data of three independent experiments, and a minimum of 3 plaques was analysed per replicate. Error bars are one standard error of the mean. p<10^-3^ using a two-tailed t-test indicated by ***. Numbers of plaques per sgRNA (-/+ L-arabinose): Control 20/15, A1 19/22, A2 17/15, A3.1 13/23, A3.2 15/13, C 16/22, 3x 16/27, 5x 11/28. Error bars are one standard error of the mean.

Surprisingly, while I found that *C* can be knocked down very efficiently at the fluorescent reporter level, I observed only a moderate effect on the time to lysis when targeting T7 RNAP during infection. Similar effects were found in a recent study that uses CRISPR-Art to target phage mRNA rather than DNA (Adler et al., 2023). Targeting T7 phage with the 3x multiplex array sgRNA showed a slight improvement over targeting *C* only, suggesting a weakly positive combinatorial effect between targeting *INTS* and *C*. The 5x multiplex sgRNA array displayed the strongest effect and a marked ∼25% increase of the time to lysis, and a relatively strong combinatorial effect of blocking several *E. coli* RNAP promoters at once (**Figure 3B**).

Finally, the efficiency of plaquing (EOP), which is a measure of infection productivity of the phage, was largely unaltered across all sgRNAs (**Figure 3C**), but there were moderate to strong effects on plaque size. Here, the order of effects followed the results of the time to lysis experiments. Blocking *A2*, *A3.1* and *A3.2* showed marked effects while blocking *A1* had the strongest effect, leading to a ∼3fold decrease in plaque size. Also, dCas9 knockdown of T7 RNAP using a single sgRNA had moderate effects on plaque size reduction, thus following the trend observed in the time to lysis experiments. Again, the 5x multiplex approach was very efficient leading to a ∼8-fold reduction in plaque size (**Figure 3D**). These results show that while dCas9 causes imperfect blockage or knockdown of T7 during infection, there is an substantial additive effect when using multiplex sgRNAs. Furthermore, while dCas9 cannot mediate resistance to T7 phage infection or affect EOP, it is capable of interfering with T7 phage during the infection process and consequently affect plaque size.

## Discussion

CRISPRi approaches based on *S. pyogenes* dCas9 have been championed in the last decade as highly flexible tools to study gene function across many different microbial species ((Qi et al., 2013; Peters et al., 2016; Liu et al., 2017; Lee et al., 2019; Peters et al., 2019; Mutalik et al., 2020; Rishi et al., 2020; Liu et al., 2021), and to construct synthetic gene regulatory circuitry (Kuo et al., 2020; Rueff et al., 2023). Until recently, there was little evidence of the usability of CRISPRi or the popular *S. pyogenes* CRISPRi-dCas9 to study phage gene function or to tune phage life cycles, although two reports suggest that phages or selected targets are amendable to *S. pyogenes* CRISPRi-dCas9 knockdown (Rousset et al., 2018; Widom et al., 2019). In this work, I explored the ability of CRISPRi-dCas9 to interfere with and tune the lifecycle of T7 phage by specifically targeting *E. coli* RNAP-dependent promoters which are essential for DNA internalisation and lifecycle execution (Calendar, 2006). By using a small set of sgRNAs and two multiplex constructs, this work is a proof-of-concept study probing the ability of dCas9 to interfere with the lifecycle of T7 phage.

By building and using a set of T7 phage fluorescent reporters, this study builds a tool to measure the strength of *E. coli* RNAP promoters annotated on the T7 genome and establishes a clear order of strength across these promoters. Promoters *C* and the promoter array within *INTS* are exceptionally strong, and the promoter array is nearly three times stronger than *C*. The strength of the minor promoters *B* and *E* was comparable to IPTG-induced P_LlacO1_, which is a synthetic promoter based on the strong phage Lambda P_L_ promoter (Lutz and Bujard, 1997). Furthermore, *INTS* has a substantial detrimental effect on growth **(SI Figure 2)** that stems either from unconstrained expression of YFP, or expression of an *ocr* fragment that is included in the fluorescent reporter construct and that may be otherwise toxic at high expression levels. The stabilised *INTS* reporter displayed a 45bp deletion upstream of the *ocr* fragment which truncates the 5’ end of the 0.3 mRNA fragment including the putative RBS of the *ocr* fragment, suggesting that this fragment is in fact detrimental to growth. The stabilised *INTS* reporter plasmid also had a measurable detrimental effect on growth which could be based on YFP overexpression or excess titration of host RNAP from the limited cellular pool **(SI Figure XY)**. Nevertheless, this raises the question whether the early steps in the infection process starting with host RNAP binding and early gene transcription leads to an immediate effect on cell growth by either titration of RNAP or by expression of toxic early gene products. A recent report studying T7 phage infection at the single-cell level doesn’t support this notion, but nevertheless shows that host cell growth stalls within the first 2-3mins of phage genome incorporation into the cell (Wedd et al., 2024).

Probing the promoter array within *INTS* with individual sgRNAs showed that there is no dominant promoter within this region as blocking *A1* had very weak or no effect on YFP fluorescence knockdown, suggesting that *E. coli* RNAP can initiate expression from the *A2* and *A3* promoters downstream of the roadblock. Concurrently, the strongest knockdown effects were apparent when either *A2* or *A3* were targeted, implying that initiation of transcription is less efficient downstream of *A2* or *A3*, or that dCas9 roadblock at these positions had other polar effects on RNAP access further downstream. Placing another sgRNA (A3.2) further downstream into the beginning of the *ocr* coding region had no effect at all. One explanation is that the knockdown efficacy of dCas9 is inversely proportional to promoter strength (knockdown efficacy decreases with increasing promoter strength), a factor that may explain the low efficacy of targeting *INTS* in general. Another possibility is that sgRNAs A2, A3.1 and A3.2 were less efficient by design and binding location within *INTS*. Another factor that could lower CRISPRi-dCas9 efficacy (which has been reported to be at 90-99% knockdown efficiency (Qi et al., 2013; Peters et al., 2019)) is that the fluorescence knockdown targets are located on an oriSC101 plasmid that has about 4 copies/cell (Jahn et al., 2016), although this would require either sgRNAs or dCas9 abundance to be limiting. Other studies reporting higher efficacies have targeted single-copy and chromosomally inserted targets (Qi et al., 2013; Peters et al., 2019), which was not applicable for the work presented here.

The results also show that the *E. coli* RNAP promoter *C* in front of the T7 RNAP is very strong, but that it can be knocked down with relatively high efficiency. It can also be speculated that, due to its high strength, promoter *C* could act together with the internalisation signal to facilitate internalisation, or alternatively aid internalisation of mutant phages lacking *A1*, *A2*, and *A3* (Garcia and Molineux, 1996). In turn, promoter *B* which has been implicated as accessory promoter for internalisation is ∼3x weaker than *C* and only mildly stronger than P_LlacO1_.

Transcriptional fluorescent reporters have been extensively used to study differential gene regulation and transcription in *E. coli* or *Salmonella typhimurium* (Zaslaver et al., 2006; Freed et al., 2008; Silander et al., 2012). Similar studies have not been conducted using phage-derived *E. coli* promoters, likely due to the unavailability of an appropriate phage *E. coli* promoter collection. The promoter plasmids constructed for the work presented here could be used in future studies that explore their activity in different environments. This could include testing a variety of carbon sources and growth conditions, different ecological contexts such as mixed species cultures or biofilms that may influence promoter expression and subsequently affect the lifecycle and infectivity of T7 phage under these likely more natural conditions.

Surprisingly, the results presented here show that blocking the first strong promoter *A1* on *INTS* had strong effects on the time to lysis and infectivity of the phage, but not the expression of the fluorescent reporter construct. This suggests that the chronological appearance of T7 phage DNA in the cell affects the efficacy of CRISPRi-dCas9 towards T7. Nevertheless, this result is surprising as a suite of elegant experiments has established that the internalisation signal is ejected independent of *E. coli* or T7 RNAP activity into the cell, and that it was thought to be initially inaccessible to host proteins (Moffatt and Studier, 1988; Garcia and Molineux, 1995; 1996). This is based on the observation that T7 DNA is not degraded by host restriction endonucleases during the first few minutes after appearing withing the cell (Moffatt and Studier, 1988). Also, a GATC motif located in the T7 early region that is supposed to be rapidly methylated by DNA adenine methyltransferase (Dam) upon T7 genome entry remains unmethylated for extended time spans (Garcia and Molineux, 1995). One possible explanation for the results shown in this work here is that sequences upstream of *A1* are required to load *E. coli* RNAP and blocking *A1* interferes with this process, or that *A1* plays a dominant role during the RNAP-dependent phase of the T7 genome internalisation process that can’t be fully compensated for by *A2* or *A3*. Another possibility is that A2, A3.1 and A3.2 sgRNAs are simply less efficient in interfering with T7 DNA internalisation based on design, although A2 and A3.1 were able to knock down expression of the fluorescent reporter. Previous work has shown that sgRNAs targeting the same genomic feature can have varying effects (Rishi et al., 2020), but the reasons for sgRNA-based variability in knockdown efficiency, other than effects caused by the “bad seed” effect (Rostain et al., 2023), are not very well understood (Cui et al., 2018).

Two recent reports have now shown that both temperate phages λ and P1 are amenable to knockdown by dCas12a (Piya et al., 2023) and phages T4 and T7 to dRfxCas13d or CRISPRi-ART (Adler et al., 2023). While dCas12a has somewhat limited target range due to its relatively large and T-rich PAM (TTTV), dRfxCas13d targets transcripts and may find broad use as tool to interrogate both phage and bacterial gene function. Overall, although *S. pyogenes* CRISPRi-dCas9 is less efficient than dCas12a, it offers greater flexibility due to its NGG PAM and usability to study knock-down effects and combinatorial effects, even if it doesn’t completely abolish T7 infectivity. Also, previous studies that have aimed at controlling the lifecycle of T7 through an “infectivity switch” rely on phage engineering (Chitboonthavisuk et al., 2022). The work presented here shows that similar, albeit milder effects on the phages time to lysis, can be achieved by rational programming of dCas9 with sgRNAs, opening new possibilities for rational engineering and programming of otherwise lytic bacteriophages for synthetic biology applications.

## Acknowledgements

I thank Remy Chait for supplying T7 phage stock, Calin Guet for pZS and pZA plasmids, Alice Thurston for very early work on the project, Ben Temperton for comments on the manuscript, and the “Bacterial Pathogenesis Group” at the University of Exeter for insightful and stimulating discussions. This work was funded by University of Exeter start-up funding to TB.

## Material and Methods

### Growth media and chemicals

Cloning and standard molecular biology techniques using strains outlined below was done in LB broth (Melford, 10 g/l NaCl) and LB agar (prepared from LB broth and 1.5g/L agar, Oxoid LP0011), and antibiotics were added as indicated. Calcium chloride dihydrate (CaCl_2_, Sigma, C3306), Magnesium sulfate (MgSO_4_, Sigma, M2643) were prepared as 1M stock solutions and sterilised by autoclaving. Chloramphenicol (Sigma C0378) and Ampicillin (Sigma A9518) were prepared as 15mg/ml and 100mg/ml stock solutions in 100% ethanol and milipore water, respectively. The latter was filter-sterilised, and aliquoted stocks were stored at −20°C. L-arabinose (Sigma A3256) was prepared as a 1M (15%) stock solution in Millipore water and sterile-filtered. Isopropyl beta-D-1-thiogalactopyranoside (IPTG, Sigma 16758) was dissolved to 1M concentration in milipore water, sterile filtered and stored in small aliquots at −20°C. T7 phage stock was generously supplied by Dr. Remy Chait, University of Exeter, and was stored at 4°C in sterile-filtered SM buffer (200mM NaCl_2_, 10mM MgSO_4_, 50mM Tris-HCl pH7.5).

### Strains, plasmids and enzymes used in this study

Fluorescence knockdown experiments, phage lysis and plaquing experiments were done using TB549, a BW25113 derivative (F^-^ LAM^-^ *rrnB3* DE*lacZ*4787 *hsdR*514 DE(*araBAD*)567 DE(*rhaBAD*)568 *rph-1*) with a single-copy insertion of Para-dCas9 into *att*HK022. Promoter strength comparisons in the absence of dCas9 and sgRNAs were done in BW25113, respectively. Strains were grown at 37°C for standard molecular methodologies and for fluorescence knockdown experiments. T7 time to lysis experiments were conducted at 30°C, and plaque size and EOP experiments at 22°C. Strains were cryopreserved at −80°C in 15% sterile glycerol, and cultures were grown from freshly streaked individual colonies on agar plates.

Plasmid construction was conducted using the isothermal Gibson Assembly method and New England Biolabs (NEB) Gibson Assembly Master Mix (#E2611). Assemblies were transformed into commercially available and chemically competent cells, NEB5α, following the manufacturers guidelines. pAH68-frt-cat-P_araBAD_-dCas9, an R6K ori plasmid (Haldimann and Wanner, 2001; Bergmiller et al., 2017) was constructed using the pir+ strain DH5αpir. Inserts of all assembled plasmids were verified using Sanger sequencing or whole plasmid sequencing using Oxford Nanopore technology (ONT, Eurofins, Germany). pZA3 (ChlorR) and pZS12-YFP Venus (AmpR) plasmids with p15a and SC101 origins of replication (Lutz and Bujard, 1997) were a gift from Calin Guet (Institute of Science and Technology, Klosterneuburg, Austria).

### Strain and plasmid construction

To construct pAH68frtcat-P_araBAD_-dCas9 and TB549, P_araBAD_-dCas9 was amplified from pZA3-P_araBAD_-dCas9 (Bergmiller lab strain collection) with primer pair dCas9 FW and dCas9 RW and assembled with a PCR product of pAH68-frt-cat that was amplified with primers pAH68frt-cat FW and pAH68frt-cat RW. dCas9 was originally amplified from plasmid pJMP1159 (supplied by Addgene) removing the myc tag (Peters et al., 2019). The sequence of P_araBAD_ was from pLA2 (Haldimann and Wanner, 2001) which has its own native ribosome binding site. The sequence of the resulting plasmid pAH68-frt-cat-P_araBAD_-dCas9 was verified by whole-plasmid sequencing using ONT (Eurofins, Germany). pAH68frtcat-P_araBAD_-dCas9 was then inserted into *att*HK022 of BW25113 using pAH69 and its single-copy status was verified by PCR following previously described methods (Haldimann and Wanner, 2001). The Chloramphenicol resistance marker was subsequently cured using the helper plasmid pCP20 (Cherepanov and Wackernagel, 1995), resulting in the AmpS ChlorS strain TB549.

Fluorescence reporter plasmids were constructed by generating a PCR product of pZS12-YFP Venus removing P_LlacO1_ using primers pZS1 FW and pZS1 RW. T7 phage fragments were synthesized as gBlocks by Integrated DNA Technologies (IDT, Belgium) with overlapping matching ends and Gibson-assembled with the pZS12-YFP Venus PCR product. All cloned T7 DNA fragments are listed in **Table 1**. Briefly, the *INTS* fragment spanned promoters *A1*, *A2*, *A3* and a short fragment of *ocr* with an in-frame stop codon. All other T7 DNA fragments included upstream and downstream sequences, and all fragments of coding regions were designed to have an in-frame stop codon. These fragments were then placed in front of the ribosome binding site of YFP Venus using Gibson assembly, and all inserts were verified by Sanger sequencing using primer pZS1-AmpR-seq.

**Table 1.**
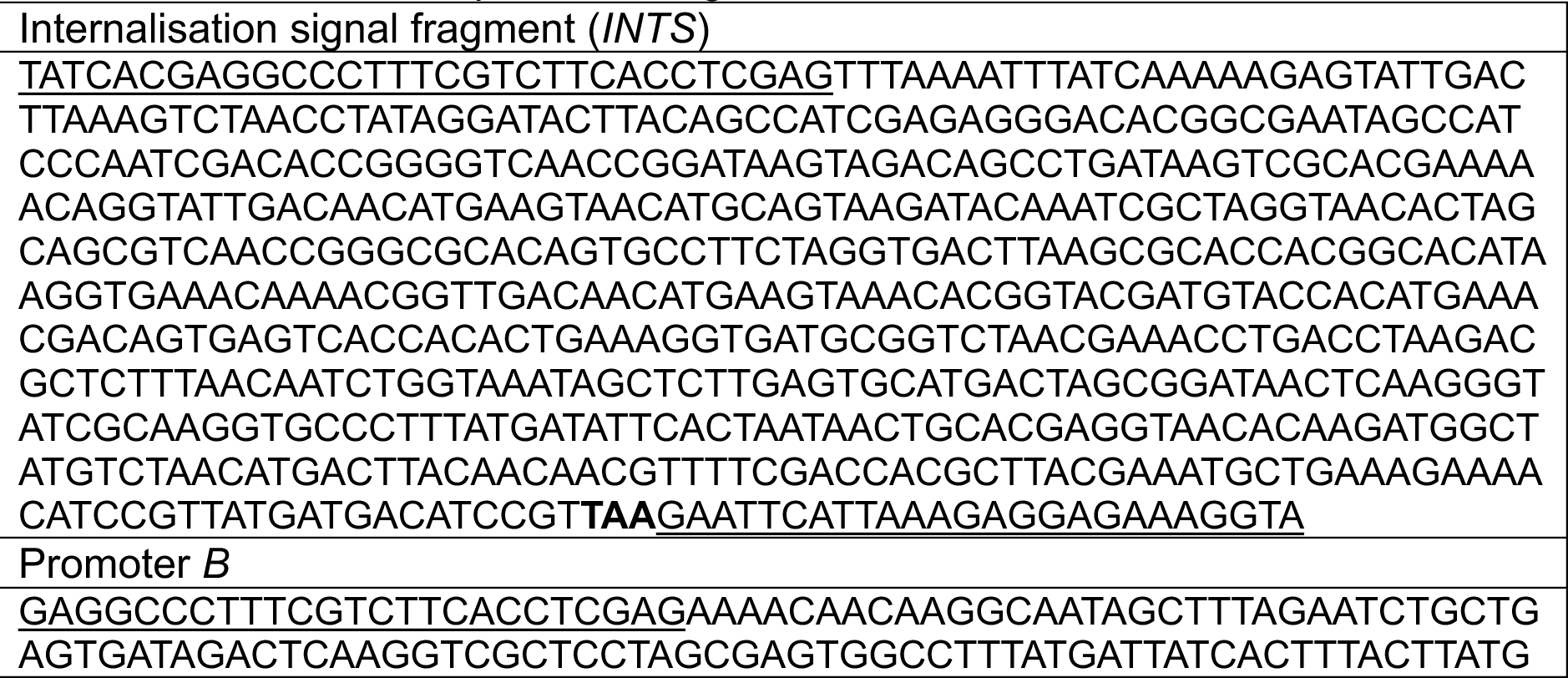
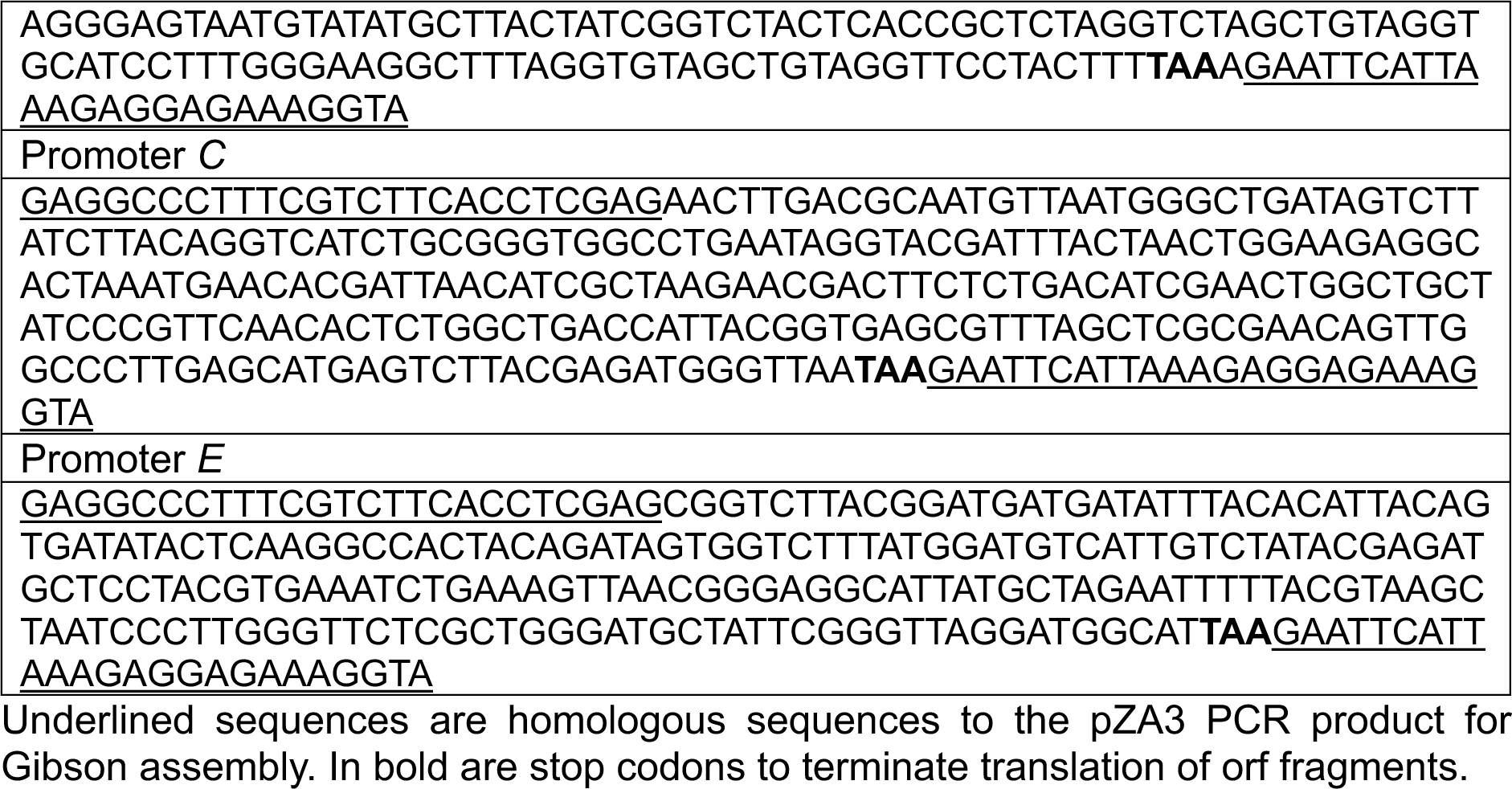
T7 *E. coli* RNAP promoter fragments.

All sgRNAs were designed following principled described in (Banta et al., 2020) and cloned into pZA3. Briefly, sgRNA sequences were selected either by identifying PAMs within the non-coding *INTS* region and designing sgRNA such that they would target the non-template strand, or by using a Python script (Banta et al., 2020) for the coding regions of gp1 (T7 RNAP) and gp0.5. All sgRNA sequences are listed in **Table 2**. The constructs were then designed such that sgRNAs were expressed by a constitutive version of P_trc_ lacking lacO sites (Banta et al., 2020) followed by a “handle” sequence and a strong transcriptional terminator. sgRNAs A3.1, A3.2 and C were designed as gBlocks with overlapping matching ends to the pZA3 PCR product generated with primers pZA3 FW and pZA3 RW and terminators Lt3, T1 and T0. sgRNAs A1, A2, and the Control sgRNA (a random 20bp DNA sequence with 50% GC content) were then constructed by using the promoter C sgRNA plasmid pZA3-C as PCR template (amplified with pZA3-C FW and pZA3-C RW) and by assembling pZA3-A1, pZA3-A2, and pZA3-Control by using single stranded oligonucleotides that had matching ends to the pZA3-C PCR product.

**Table 2.**
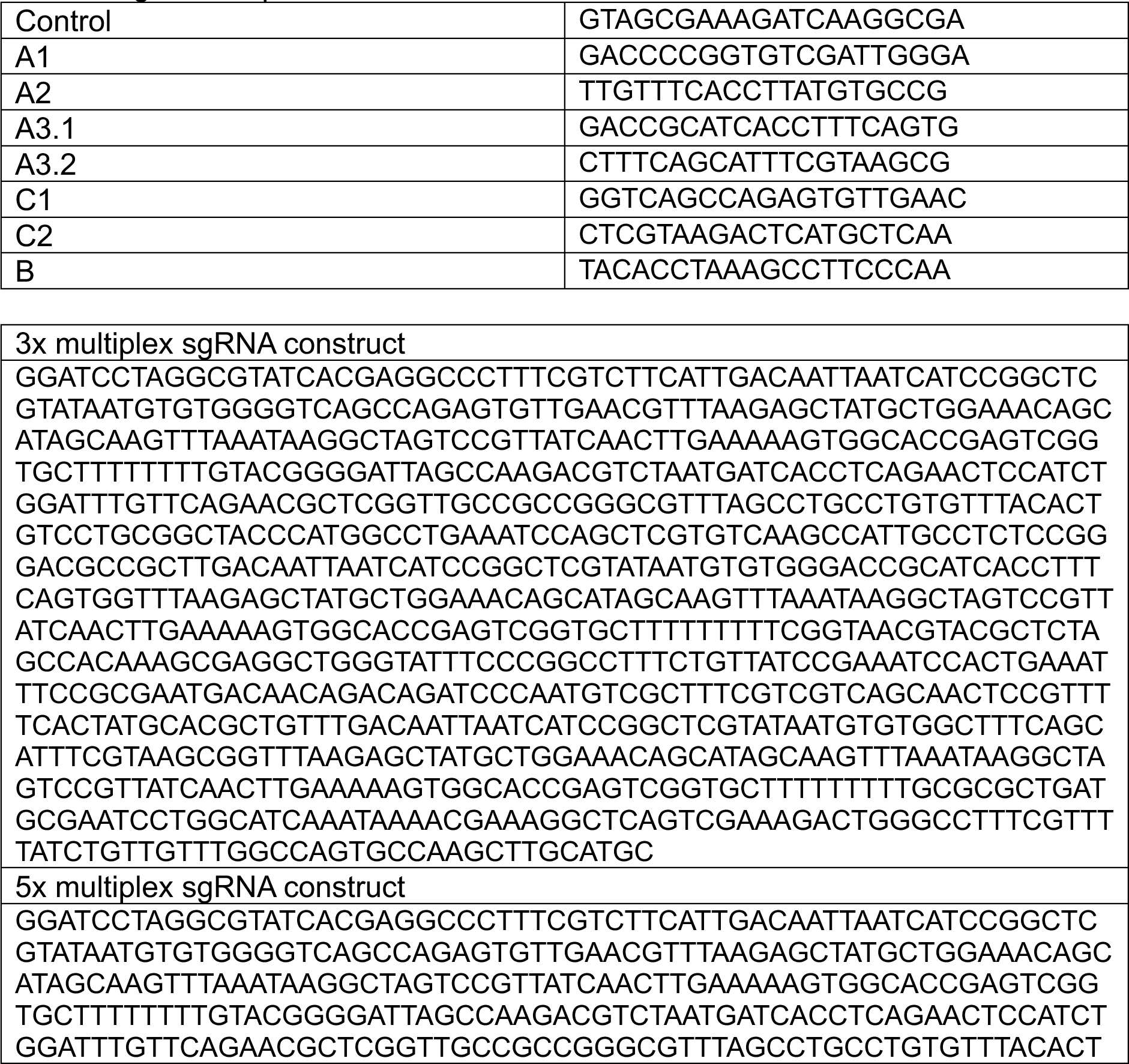
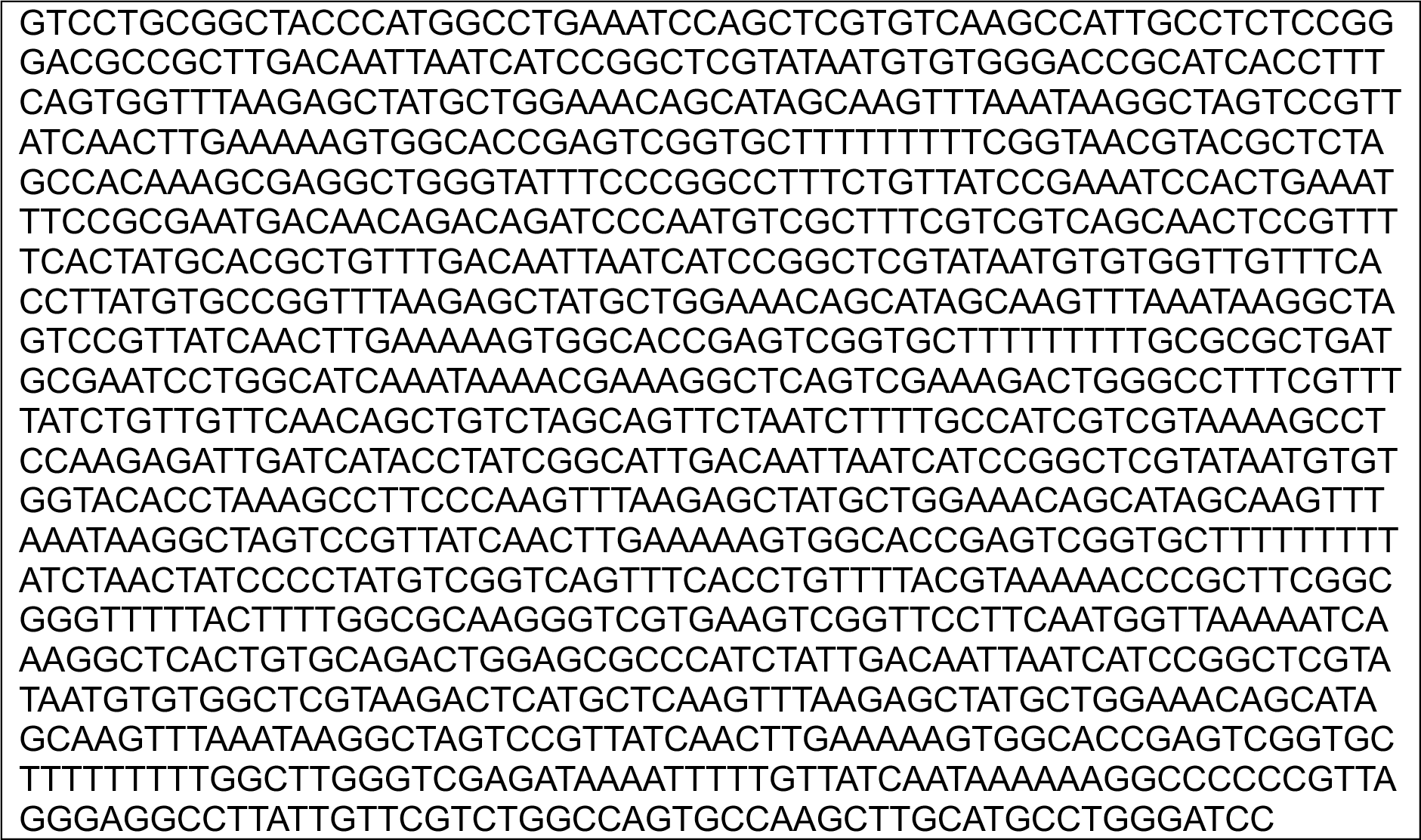
sgRNA sequences.

**Table 3.**
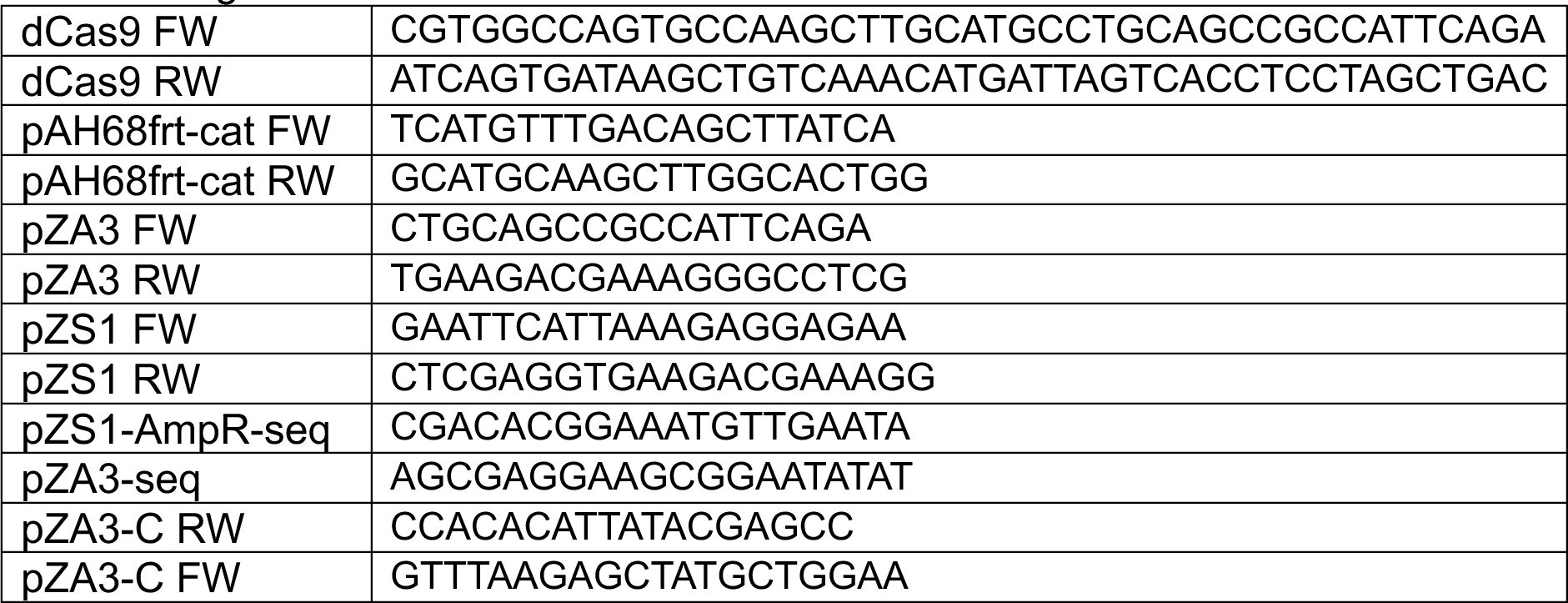
Oligonucleotides.

**Table 4.**
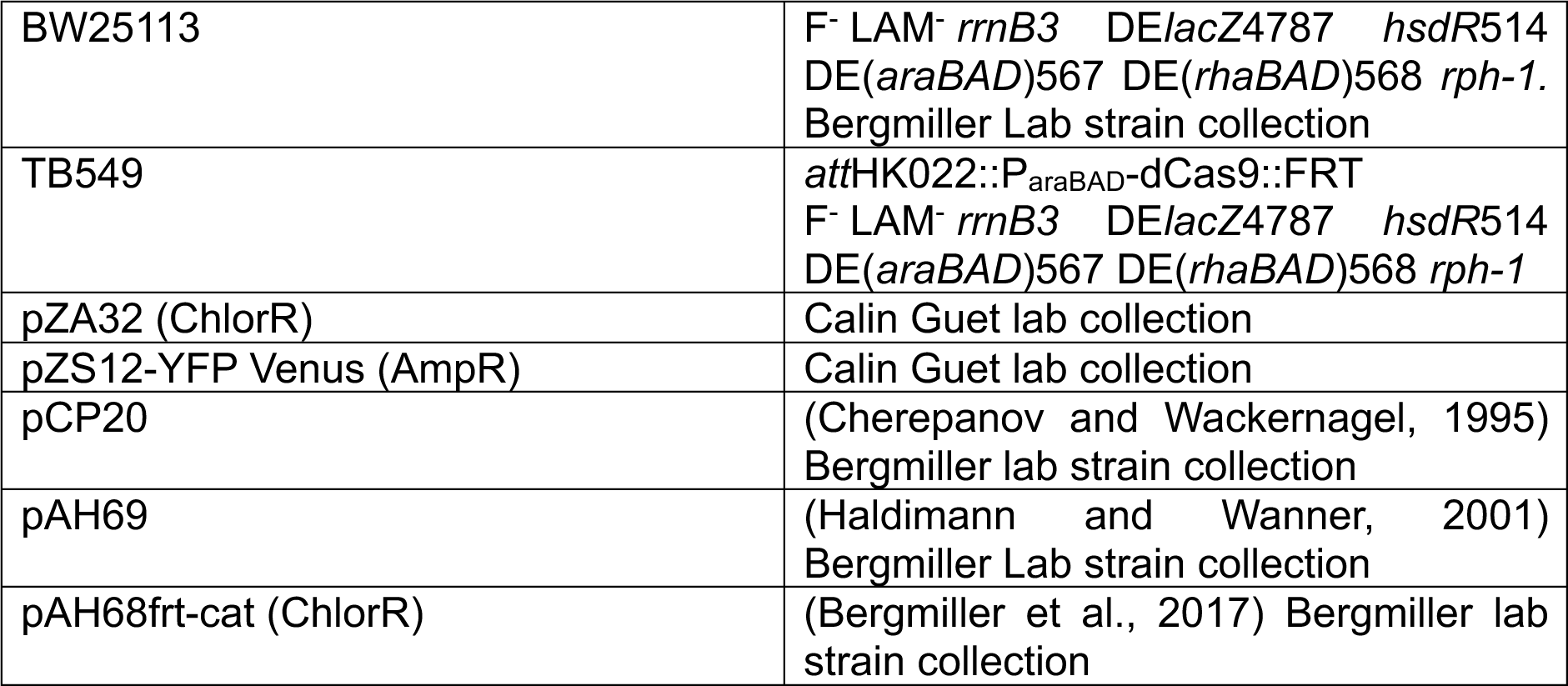
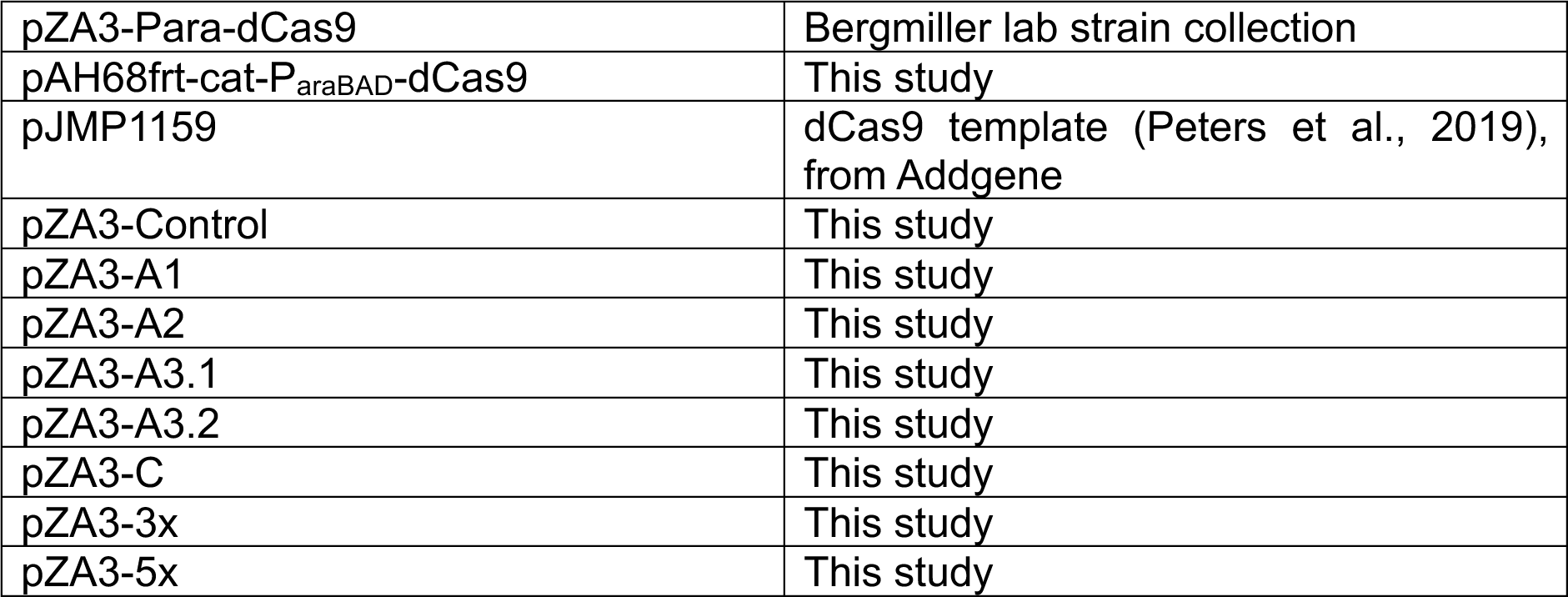
Plasmids and strains.

|  |  |
| --- | --- |
| BW25113 | F<sup>-</sup> LAM<sup>-</sup> *rrnB3* DE*lacZ*4787 *hsdR*514<br>DE(*araBAD*)567 DE(*rhaBAD*)568 *rph-1*.<br>Bergmiller Lab strain collection |
| TB549 | *attHK022::P<sub>araBAD</sub>-dCas9::FRT*<br>F<sup>-</sup> LAM<sup>-</sup> *rrnB3* DE*lacZ*4787 *hsdR*514<br>DE(*araBAD*)567 DE(*rhaBAD*)568 *rph-1* |
| pZA32 (ChlorR) | Calin Guet lab collection |
| pZS12-YFP Venus (AmpR) | Calin Guet lab collection |
| pCP20 | (Cherepanov and Wackernagel, 1995)<br>Bergmiller lab strain collection |
| pAH69 | (Haldimann and Wanner, 2001)<br>Bergmiller Lab strain collection |
| pAH68frt-cat (ChlorR) | (Bergmiller et al., 2017) Bergmiller lab<br>strain collection |
| pZA3-Para-dCas9 | Bergmiller lab strain collection |
| pAH68frt-cat-P<sub>araBAD</sub>-dCas9 | This study |
| pJMP1159 | dCas9 template (Peters et al., 2019), from Addgene |
| pZA3-Control | This study |
| pZA3-A1 | This study |
| pZA3-A2 | This study |
| pZA3-A3.1 | This study |
| pZA3-A3.2 | This study |
| pZA3-C | This study |
| pZA3-3x | This study |
| pZA3-5x | This study |

3x and 5x multiplex sgRNA constructs were synthesised as full-size constructs by GenScript (GenScript Biotech, UK), cut out from the delivered plasmid using BamHI, and Gibson-assembled into pZA3. Multiplex constructs were designed such that each P_trc_-sgRNA-terminator module was preceded by 75bp random DNA sequence and either 3 or 5 different terminator sequences were chosen to minimise sequence redundancy. For 3x, terminators were T0-Lt3-T1, and for 5x T0-Lt3-T1-garPLRK-L3S1P13.pZA3 plasmids were all verified by Sanger sequencing using primer pZA3-seq.

All plasmid DNA sequences are available as GenBank files in the supplementary files.

### Growth rate experiments

To estimate their doubling time, biological duplicates of bacterial strains were grown overnight to saturation, and if required, antibiotics added to select for plasmids. Overnight cultures were then split into three technical replicates and diluted 1:200 into fresh broth in transparent 96well microtiter plates (Greiner #655161), again with fresh antibiotics added to select for plasmids, and plates were sealed using clear adhesive film (Fisher Scientific #15963620). The plates were then incubated at 37°C and optical density at 600nm measured every 5 minutes in a BMG Clariostar Plus. Before each measurement, the plate was shaken double-orbitally at 600rpm for 10sec. Growth rates and doubling times were then estimated using a custom-made R script that fits a linear slope to the steepest part of log-transformed data over 5 timepoints using a R^2^ criterium of 0.98. Two separate doubling times for the initial *INTS* fluorescent reporter plasmid were estimated by fitting curves over each 5 timepoints to the two separate growth phases shown in **SI Figure 2**. Growth rates of individual promoter constructs in the absence of sgRNA plasmids was done in biological quadruplicates, while TB549 with or without sgRNA plasmids and dCas9 induction was done in biological duplicates with technical triplicates.

### Fluorescent reporter strength and knockdown experiments

Fluorescent knockdown experiments of reporter plasmids were done in TB549. To do so, TB549 was sequentially transformed with matching pZA3-sgRNA plasmids or the pZA3-Control plasmid and pZS1 reporter plasmids. Overnight cultures of double transformants were inoculated 1:200 into fresh LB broth with antibiotics in black 96well microtiter plates with clear bottom (Greiner #655090), sealed with clear adhesive film and placed in a temperature-controlled BMG Clariostar Plus plate reader at 37°C. OD_600nm_ and YFP fluorescence reads (515nm excitation, 530nm emission) using the enhanced sensitivity range were taken every 5mins that were preceded by 10sec of shaking in a double-orbital pattern at 600rpm. After subtracting the blank values of control wells, fluorescence data was normalised by OD_600nm_ (Fluorescence/OD_600nm_). Promoter strength was measured of each 4 independent biological replicates, and uninduced pZS12-YFP Venus used to normalise promoter strength data was done in biological triplicates. To compare promoter strength across fluorescent reporters in the absence of sgRNA plasmids, single time points at 100mins were analysed and normalised to the mean fluorescence of TB549 harbouring uninduced pZS12-YFP Venus.

Fluorescent knockdown experiments were done in biological duplicates that were split into two sets of technical triplicates into which either no L-arabinose (controls, no dCas9 expression) or 0.1% L-arabinose was added (dCas9 expression and fluorescence knockdown), yielding a total of 6 control cultures and 6 fluorescence knockdown cultures for each fluorescent reporter-sgRNA combination. To analyse the knockdown efficiency of fluorescent reporters, data was normalised to the mean of respective controls lacking arabinose. Data from timepoint 400mins (early stationary phase) was analysed for *INTS* and promoter *C* knockdown.

The choice of timepoint for quantifying promoter strength or knockdown efficiency affects the values that are used to quantify these parameters. Overall, the timepoints were chosen such that fluorescence levels had stabilised or plateaued, which occurs during early stationary phase. This is likely driven by dilution of highly expressed and otherwise proteolytically stable YFP upon dCas9 knockdown. For some of the data, in particular for the *INTS*-YFP construct, there is variation arising in fluorescent levels into later stationary phase which could be caused by cell death or other unknown factors. Promoter strength experiments are less affected by equilibrating YFP levels as no knockdown occurs, and thus a robust time point to assess fluorescent levels is mid-to late exponential phase assuming near steady-state growth and an equilibrium between YFP production and dilution at this growth phase.

To compare fluorescent reporters and knockdown efficiencies on plates which serve as visual guides to compare knockdown efficiencies, agar plates with antibiotics and with or without 0.1% L-arabinose were sectored, and individual colonies replica-streaked onto separate sectors. Images of plates were taken using a custom-made fluorescent macroscopic imager (“macroscope” (Chait et al., 2010)) and their brightness and contrast adjusted on a linear scale using ImageJ.

### T7 phage time to lysis experiments and data analysis

All T7 phage lysis experiments to determine the time to lysis were done in transparent 96-well microtiter plates (Greiner #655161) in a BMG Clariostar Plus plate reader at 30°C. Biological duplicates of overnight cultures were diluted 1:100 into fresh LB with 0.1mM CaCl2 and 10mM MgSO4 and grown with or without 0.1% L-arabinose for 2.5h reaching OD600nm of 1. Then, a T7 phage dilution series was made in a 96well plate containing columns of LB with 0.1mM CaCl2 and 10mM MgSO4 and with or without 0.1% arabinose, cultures split into technical triplicates, and cells added such that they were diluted 1:10 into wells with three different phage dilutions (10^-8^, 10^-9^ and 10^-10^ PFU). OD600nm measurements were done every 5 minutes and preceded by a 10sec 600rpm double-orbital shake for 300mins. To find the point of culture lysis, the inflection points at which the slope turns negative were determined by fitting a slope using a sliding window of 4 time points to OD_600nm_ growth curves. The interception point of slopes with the x-axis was then determined using the Excel function “Intercept”. The onset of culture lysis was found to be increasingly variable at higher dilutions of 10^-9^ and 10^-10^ PFU due to extremely low phage numbers and MOI, and subsequently data from wells with 10^-8^ PFU was analysed (equalling an MOI of 10^-5^). The change in time to lysis was then calculated by normalising the time to lysis of cultures grown with 0.1% L-arabinose to the mean time to lysis of control cultures lacking L-arabinose.

### Top agar experiments

Efficiency of plaquing (EOP) and plaque size experiments were done using the spot dilution method (Adams, 1959). Briefly, top agar (0.5% LB agar) was stored in 5ml aliquots in a 50°C water bath with 0.1mM CaCl2, 10mM MgSO4, 15µg/ml Chloramphenicol and with or without 0.1% L-arabinose. Strains carrying sgRNA plasmids were cultured with or without 0.1% arabinose for 3.5h, and then 1ml of bacterial culture was added to the top agar, the tube inverted, its content poured on top of LB agar plates, and allowed to dry. 3µl of phage dilution were spotted onto the top agar, allowed to dry, and plates incubated at 22°C for 20h. Images of plates were taken using a BioRad GelDock imager, and plaque area was measured using the oval selection tool in ImageJ. Three independent biological replicates were done and at least three plaques of each plate measured.

---

**Supplementary Information Figure 1.**
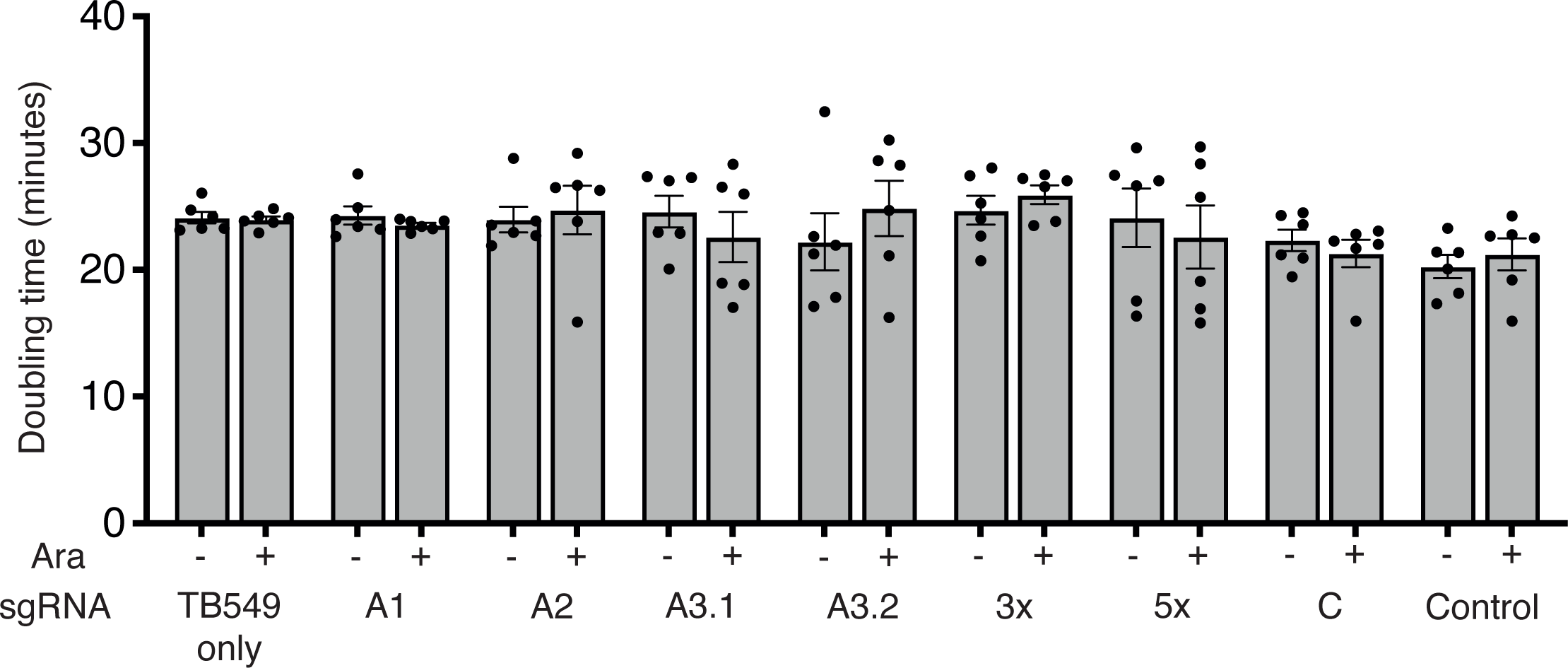
Growth rates of TB549 expressing sgRNAs in the presence or absence of dCas9 induction. Shown are doubling times in minutes of TB549 expressing sgRNAs in the absence or presence of dCas9 induced with 0.1% L-arabinose. There is no statistical significance between uninduced and induced dCas9 expression for all sgRNAs, or TB549 alone (two-tailed t-tests). N=6 for all bars shown. Error bars are one standard error of the mean.

**Supplementary Information Figure 2.**
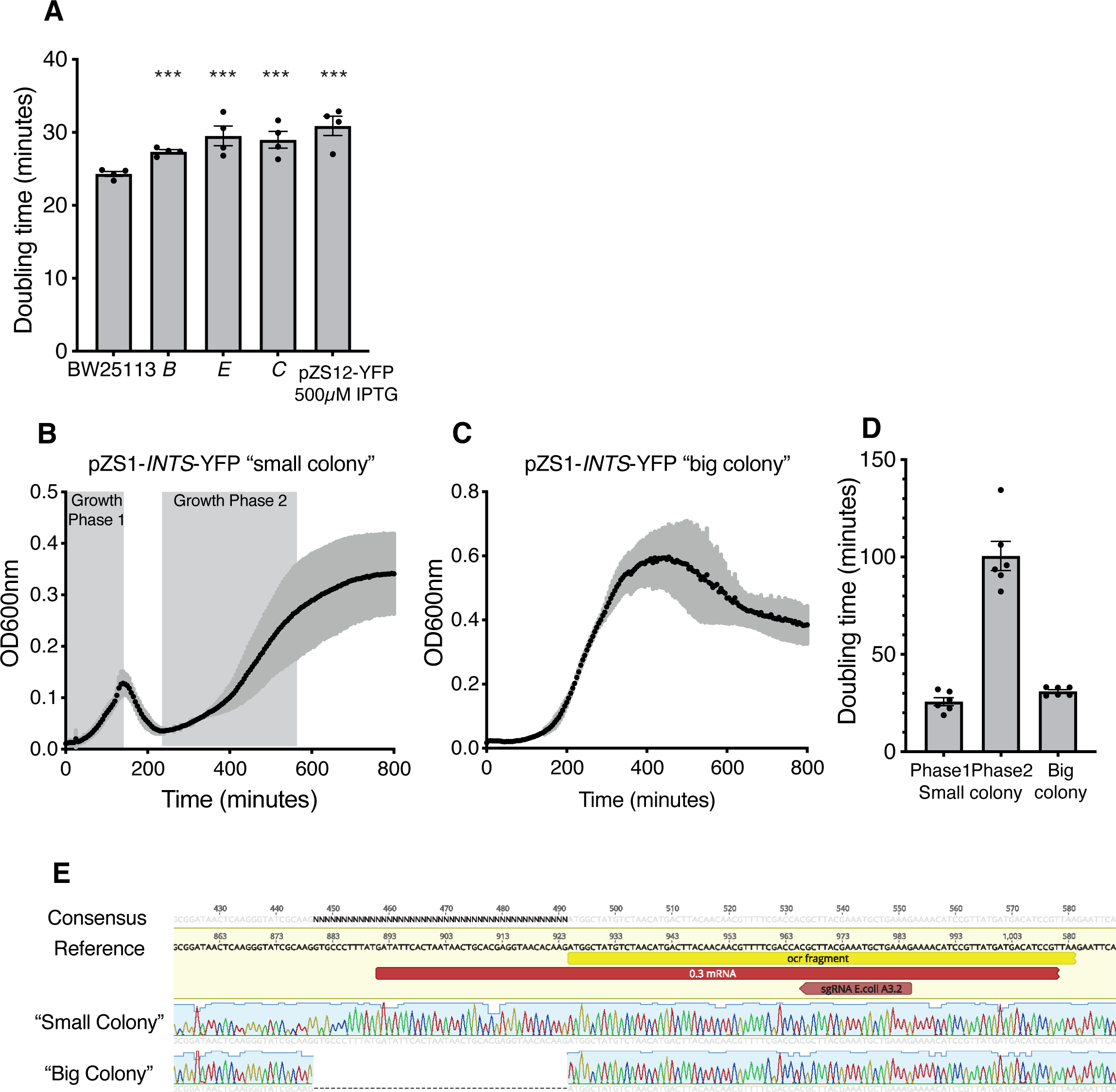
Growth rates of strains expressing fluorescent promoter reporters. A) Doubling times of BW25113 expressing respective fluorescent reporter plasmids or pZS12-YFP Venus induced with 500µM IPTG. All fluorescent reporters increase doubling time compared to BW25113 (pairwise t-test p<10^-3^ indicated by ***). N=4 for all data shown. B) Growth curve of the pZS1-*INTS*-YFP “small colony” variant in TB549 in the absence of dCas9 expression. Indicated are the two growth phases that appear to be separated by partial culture lysis. C) Growth curve of the pZS1-*INTS*-YFP “big colony” variant. D) Doubling times of growth phases 1 and 2 of pZS1-*INTS*-YFP “small colony” and pZS1-*INTS*-YFP “big colony” variants. E) Sanger sequencing chromatograms showing a 45bp deletion in the pZS1-*INTS*-YFP “big colony” reporter plasmid and the original “small colony” variant sequence. N=6 for growth curves. Bold line shows the mean, shaded curve is one standard error of the mean. Error bars are one standard error of the mean.

